# The ear receives only weak parametric information about saccade directions

**DOI:** 10.64898/2026.11.24.768848

**Authors:** Nancy Sotero Silva, Christoph Kayser

**Affiliations:** Cognitive Neuroscience, Faculty of Biology, Bielefeld University, Bielefeld, Nordrhein-Westfalen, Germany

**Author notes:** Correspondence should be addressed to: NSS.

**Keywords:** eardrum oscillations, saccades, oculomotor control, hearing

## Abstract

Eye movement-related eardrum oscillations (EMREOs) are acoustic signals recorded in the ear canal induced by saccades, whose amplitude and time course relate to the saccade vector. While horizontal saccades evoke robust, and largely opposing EMREO signals in the two ears, the reliability and informational content of vertical EMREOs has remained more elusive. Here, we analyzed bilateral EMREOs in human participants performing visually guided horizontal and vertical saccades. Consistent with previous work, horizontal saccades induced stereotyped, opposing waveforms across ears. In contrast, vertical EMREOs exhibited distinct temporal profiles between upward and downward directions, weaker signal modulation, and substantially higher inter-individual variability, particularly for downward saccades. To quantify the parametric information carried by single-trial bilateral EMREOs about the saccade vector, we applied multivariate classification and reconstruction analyses. The cardinal saccade direction was classified above chance, with significantly higher accuracy for horizontal than vertical saccades. Similarly, the reconstruction recovered horizontal saccade components more accurately than vertical ones, yet severely underestimated saccade amplitudes with high variability. These findings demonstrate that EMREOs carry reliable categorical saccade information, but primarily along the horizontal axis, and provide only weak and noisy parametric estimates of the full two-dimensional saccade vector. They also suggest that EMREOs may be particularly driven by the differential engagement of oculomotor neuromuscular pathways across the two sides of the brainstem.

## Introduction

During saccadic eye movements the ears emit recordable signals whose sign and amplitude carry information about the saccade direction. These eye movement-related eardrum oscillations (EMREOs) are induced by oculomotor-related feedback relayed to the middle ear and possibly the cochlea (Gruters et al, 2018; Bröhl and Kayser, 2023; King et al., 2023; Lovich et al., 2023a). Recent studies show that EMREOs emerge for saccades to sensory stimuli but also with eyes closed (Sotero Silva et al., 2025; Abbasi et al., 2025; Leon et al., 2025). Furthermore, EMREOs are not affected by contralateral acoustic suppression, tilting the head, and hence mostly reflect oculomotor planning and execution (Sotero Silva, Bröhl and Kayser, 2026; Sotero Silva and Kayser, 2026). Consistently across studies, EMREOs induced by left- and rightwards saccades are opposing each other but are highly consistent across participants. Those induced by vertical saccades have been studied less frequently but were reported as more variable (King et al., 2023, Lovich et al 2023b). Some studies speculated that EMREOs may reflect direct parametric information about the saccade vector, which could in principle allow including visuo-spatial information directly when encoding sounds in the ear (Gruters et al., 2018; King et al., 2023; Lovich et al 2023a, b).

Horizontal and vertical saccades rely on distinct neuromuscular pathways to coordinate direction-specific gaze shifts. Horizontal saccades are driven by the medial and lateral rectus, which are differentially activated in both eyes (Rucker, 2010; Leigh and Zee, 2015). These two muscles are innervated by distinct cranial nerves (CN): the medial rectus by CN III and the lateral rectus by CN VI, the action of which is coordinated across the midline by the paramedian pontine reticular formation and the medial longitudinal fasciculus (MLF) (Fite and Walker et al., 1990; Pierrot-Deseilligny, 2011). It has been speculated that the opposition of the engaged neuromuscular pathways may contribute to the opposing EMREOs induced in the two ears (Sotero Silva, Kayser and Bröhl, 2025). In contrast, vertical saccades require the simultaneous coordination of four muscles per eye: upwards saccades are induced by the concurrent contraction of the superior rectus and the inferior oblique, both innervated by CN III and coordinated bilaterally via the rostral interstitial nucleus of the MLF (Moschovakis et al., 1991; Rucker, 2010; Horn and Leigh, 2011). Downwards saccades are induced by the inferior rectus (innervated by CN III) and the superior oblique (innervated by CN IV), whose trochlear nucleus receives predominantly ipsilateral, uncrossed premotor drive from the rostral interstitial nucleus of the MLF (Bender, 1980; Horn and Leigh, 2011). Hence, downward saccades engage a more complex combination of ipsilateral and contralateral pathways compared to upward saccades (Lal and Truong, 2019). This leads to interesting predictions for EMREOs, assuming that signals relating to the activation of different CNs are relevant for inducing EMREOs.

One is that EMREOs induced by left and rightwards saccades should oppose each other, in time and amplitude. This overall anti-phase pattern of horizontal EMREOs has been generally confirmed (Gruters et al., 2018; King et al., 2023; Sotero Silva, Kayser and Bröhl, 2025), although fine-grained statistical analyses also point to idiosyncratic differences in ipsi- and contralateral EMREOs (Sotero Silva, Kayser and Bröhl, 2025). A second prediction is that EMREOS induced by vertical saccades should exhibit temporal profiles that differ between the two vertical directions and from those of horizontal EMREOs. Since upwards saccades depend exclusively on CN III, while downward saccades recruit both CN III and CN IV, the respective EMREOs should not simply oppose each other. One may speculate that downward saccades could exhibit a more complex time course driven by multiple neural pathways. In addition, vertical saccades engage different nerves than horizontal saccades, and the respective EMREOs could differ clearly from each other. Few studies have investigated vertical EMREOs in detail, but the available evidence suggests vertical EMREOs appear less reliable across participants and exhibit generally a weaker modulation of time course compared to horizontal EMREO (Bröhl and Kayser, 2023; King et al., 2023; Lovich 2023b).

We here recorded EMREOs to saccades in horizontal and vertical directions in humans. We capitalized on ideas pioneered in a previous study suggesting that EMREOs provide parametric information about oculomotor behavior (Lovich 2023b) and used two approaches to test for such parametric information. One approach used a multivariate classification (decoding) framework similar to approaches often used for the decoding of neural or neuroimaging signals to test how well one can predict the saccade direction from the bilateral EMREOs. Another approach used stimulus reconstruction to directly predict the saccade vector from the EMREOs.

## Materials and Methods

### Participants

We recruited human volunteer participants among community members at a location which will be identified if the article is published. The study was approved by the authors’ university Ethics Review Board (#2024-037). All participants gave informed consent on a written form and received a 15 Euros per hour compensation. For this study we collected data from 31 adult participants from both sexes (17 female, 14 male, age range 18-40). Inclusion criteria required no history of neurological disorders, no visual impairment (visual dpt < 2 was tolerated) and positive visual ear canal inspection. Participants reported hearing loss or middle ear impairments (i.e., otitis, blockage of the ear canal) in at least one ear were excluded from the study. For three participants the eye tracker did not reliably collect data in several blocks of the experiment and the data were not further analyzed. Hence, we analyzed EMREO recordings from 28 participants.

### Experimental setup

The data were collected in a soundproofed, dark and electrically shielded booth. Participants were seated 1 m away from a sound-permeable screen (Screen International Modigliani, 2 × 1 m) with their head on a chinrest. Visual stimuli were projected (Acer Predator Z650, Acer; 60 Hz refresh rate) onto the screen and stimulus presentation was controlled using the Psychophysics toolbox (Brainard, 1997) for MATLAB (The MathWorks, 2022) and synchronized to the recording system using transistor-transistor logic (TTL) pulses. Eye movements were recorded at 1000 Hz sampling rate from the left eye using a desktop mounted infrared-based EyeLink 1000 Plus Eye Tracker (SR Research, 2015) running Host Software vs 5.15. Eye tracking calibration and validation were performed before starting a recording block with a 5-point grid. The parameters for saccade detection in the EyeLink system were velocity threshold of 30°/s, and acceleration threshold of 8000°/s². Horizontal and vertical eye position was recorded in degrees. In all experiments the eye tracker camera was aligned with the chinrest and participants’ head. In-ear microphone recordings were performed using two Etymotic ER-10C (Etymotic Research) systems. The recorded microphone signals were pre-amplified using the ER-10C DPOAE amplifiers (Etymotic Research) with gain set to +20 dB. Signals were digitized through an ActiveTwo AD-box (BioSemi) at a sampling rate of 2048 Hz.

Prior to each experiment, we ensured data quality by performing control measurements for the in-ear microphones. First, we made an empty-room measurement during which the in-ear probes were attached to the chinrest to simulate the typical in-ear position relative to participants’ heads. Then 15 seconds of silence followed by the presentation of 20 tone stimuli (10 stimuli at 100 Hz, 10 stimuli at 200 Hz, duration 400 ms) were presented from speakers behind the screen. The same measurement was repeated with the in-ear probes inserted into the participants’ ear canals. The baseline noise spectrum and the signal amplitudes in response to these tones were used to adjust the ear-probe insertion for the experiment and to calibrate the signal levels between ears, similar to other work in the field (Sotero Silva, Kayser and Bröhl, 2025).

### Experimental paradigm

We collected EMREOs during visually-guided horizontal and vertical saccades. Horizontal and vertical saccades were tested in different blocks. The horizontal saccades task comprised saccades either from a central fixation position to a left (-12°) or right (12°) target, or back from these eccentric targets to the central position. In each trial, a fixation dot (0.2° radius, white on the gray background) was presented steadily for 650 ms at the respective start location. The dot then jumped to one of the two eccentric positions (if starting in the center), or back to the central position (if starting in the eccentricities). The fixation dot remained on the screen continuously and participants were instructed to fixate this and to follow it with the eyes as quickly as possible. Each eccentric position was presented 25 times and the central position 50 times, resulting in 100 saccades per block, with a total of 200 trials split in 2 blocks. Intertrial intervals lasted between 650 and 1000 ms (uniform). The vertical task was identical, except that targets were presented upwards (+12°) or downwards (-12°) from the central position. Each eccentric position was presented 25 times, the central position 50 times, resulting in 100 saccades per block and a total of 400 trials split in 4 blocks. We collected a larger number of trials for the vertical dimension on purpose, as quantifying the reliability of vertical EMREOs was one main objective of this study.

### Analysis of EMREOs

The in-ear recordings were epoched around the onsets of individual saccades and averaged across epochs for saccades towards each direction (left, right, up, or down). For this we first detected the onset of the largest saccade observed in a 3 s period following the jump of the fixation dot and epoched the data in 400 ms windows around these onsets (-150 ms prior to onset, + 250 ms post-onset). To avoid artifacts, we only included saccades whose amplitude was between 2° to 18°, for which the initial fixation prior to the saccade was stable and for which the entire horizontal and vertical traces of the eye position were contained in an 18° x 18° window around the central fixation point. To screen for outliers the resulting data epochs were z-scored and epochs for which the maximal z-score of the EMREO signal in either ear at any point in time exceeded z = 4 were excluded from further analysis. To visualize EMREOs we averaged the non z-scored epochs (in units of mV). This resulted in one average EMREO time course per participant, ear, and saccade direction. Since not every recorded ear exhibits a clear EMREO signal, we included only those participants in the final analysis for whom the horizontal EMREOs were visually robust and exceeded a specific signal-to-noise criterion. The SNR of the epoch-averaged EMREO was calculated as the ratio of the maximal absolute deflection in a time window of 0 to 50 ms from saccade onset to the standard deviation of the signal in the time window of -150 to 0 ms. Participants with an SNR smaller than 2 in at least one ear were excluded from subsequent analysis. We based the participant exclusion solely on the horizontal EMREO, since this is known to be robust and reliable across participants and allowed for an unbiased analysis of the vertical EMREO. Based on these criteria the final analysis was based on data from n = 20 participants, for which we analyzed EMREOs induced by 191 ± 2 horizontal saccades (mean ± sem per participant) and 344 ± 8 vertical saccades.

Group level inference on the EMREO time course was based on cluster based permutation tests that allow exploiting the smoothness of the respective signals while controlling for multiple comparisons along time; similar to approaches used for neuroimaging data (Sotero Silva and Kayser, 2026; Nichols and Holmes, 2002). First level contrasts were based on paired two-tailed t tests. The outcome of these first-level tests was thresholded at p < 0.05 and the test statistic was aggregated across neighboring significant time bins (computing the max-sum as cluster-forming statistics and requiring a minimal cluster size of 5 ms). The resulting clusters in the actual data were compared to potential clusters in 10’000 sets of surrogate data, obtained after randomly flipping the effect-sign for each participant. We report clusters with a second level p < 0.05.

### Classification and reconstruction analyses

For both analyses we extracted the EMREO time courses in both ears in windows from 0 ms to 80 ms relative to saccade onset and appended these (in time) to constitute one feature vector per saccade. This window was chosen as it captures the majority of the temporal modulation of the signal time course. By appending both EMREOs we exploited the binaural EMREO signal but made no assumption about what combination of the binaural signal (binaural difference, sum, or just a monaural signal) would provide most information about the saccade. Both classification and reconstruction analyses were computed separately for each participant, hence allowing for individual differences in what pattern of the EMREO signal carries relevant information.

Classification tested how well each of the four discrete saccade directions (left, right, up, or down) can be identified on a single trial basis. This was tested using a nearest neighbor algorithm: each single trial feature vector was compared to the average feature vector for each direction (computed across all saccades except the to-be-tested one) using the Euclidean distance. The to-be-tested feature vector was then classified as the direction to which it had the smallest distance. For each participant we summarized the classification performance in a confusion matrix, quantifying how frequent each test direction was classified as each of the four candidate directions. Since we collected and included more trials for vertical than horizontal saccades, this imbalance may effectively increase the stability of the vertical class templates compared to horizontal templates and potentially inflate the classification accuracy for vertical relative to horizontal saccades. Since we found lower accuracy for vertical saccades, this imbalance is not a concern for our conclusion.

Reconstruction tested how well the continuous saccade vector (i.e., its horizontal and vertical components) can be predicted from the single trial EMREO features. This was based on ridge regression models: two separate models were computed, one predicting the horizontal component and one the vertical component. The models exploited an L2-regularized linear regression and least-squares loss function (fitrlinear in MATLAB). To prevent overfitting and selecting the optimal regularization parameter, we performed 5-fold cross-validation across a logarithmically spaced grid of 20 regularization values ranging from 10^-8^ to 10^5^. For each component, the regularization value that minimized the cross-validated mean squared error was selected, and a final ridge regression model was refitted using this optimal hyperparameter to generate the predicted saccade vectors. Note that although the saccade targets jumped only in discrete values of 12°, the actually executed saccade span a wide range of amplitudes (c.f. Fig. 1C, 2C), which ensured that the model was trained on a range of saccade amplitudes. We ensured model validity using three post-hoc sanity checks: by assessing prediction-to-target variance ratios to rule out over-regularization; by computing the cosine similarity between horizontal and vertical weight vectors to confirm distinct decoding subspaces; and by verifying the Pearson correlation between predicted horizontal and vertical vectors. We then evaluated these models by computing the correlation between actual and predicted horizontal (vertical) saccade components for each participant, and by computing the root-mean-squared deviation between these (rmse).

**Figure 1:**
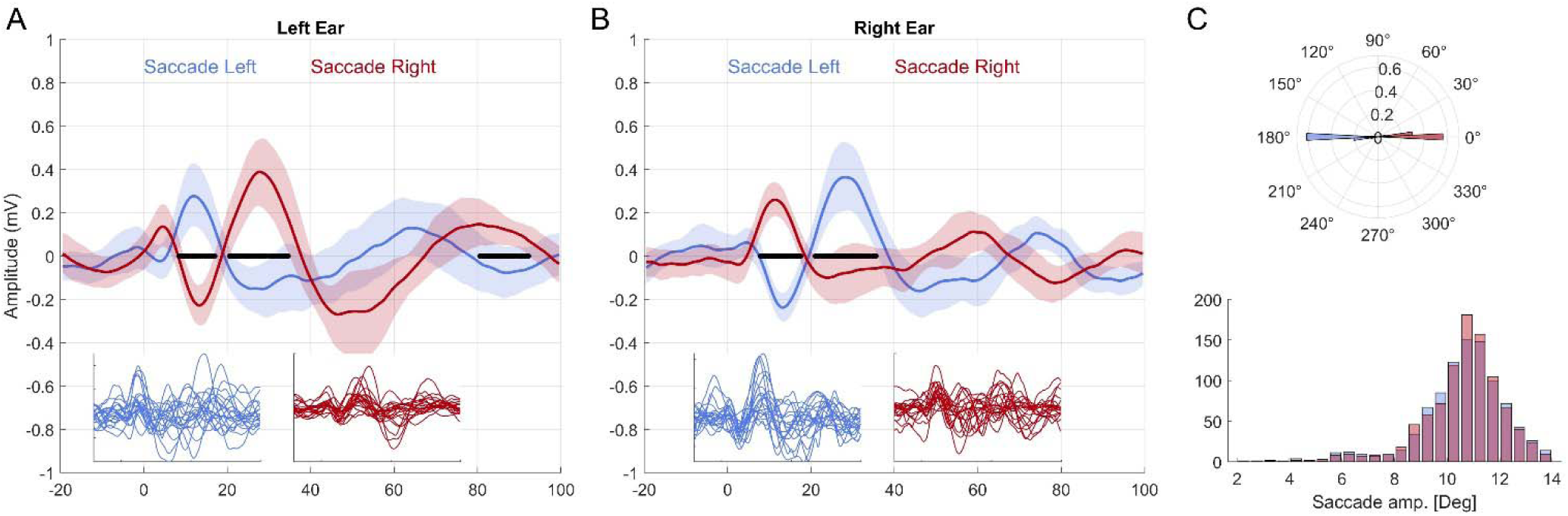
EMREOs induced by horizontal saccades. **A, B)** Group-level EMREO time course (mean and 95% percentile bootstrap confidence interval across participants) for both saccade directions in each ear. Black lines indicate time windows with significant differences (cluster-based permutation test, at least p < 0.05). Insets show the individual participant EMREOs. **C)** Distribution of saccade directions (upper) and saccade amplitudes (lower) across all included saccades and directions.

### Statistical testing

For EMREOs, statistical testing of specific null hypotheses was performed using cluster-based permutation tests described above, which control for multiple tests along time. For these we report the associated p-values for each significant cluster, the respective time window included in this, and to provide a measure of effect size, we report the associated Cohens’ D effect size of the underlying within participants t-test at the time point of maximal effect. When contrasting EMREOs between all four directions, we used the harmonic mean p-value to combine p-values across different significant clusters per ear and across ears (Wilson, 2019). This allowed us to reduce this comparison to one effective test (and p-value) per comparison of saccade directions. To compare participant-wise average saccade amplitudes between conditions, correlation values or the rmse, we used paired t-tests, for which we report Cohens’ D effect size and a Bayes factor against the null hypothesis (bf10) derived using the bayesFactorToolbox for Matlab (Krekelberg, 2024).

### Code and Data Accessibility

The code and data will be made available on github once the paper is ready for publication.

## Results

### Horizontal EMREOs

The EMREO time courses for horizontal saccades differed between saccades towards ipsi- and contralateral targets (Fig. 1A, B), in line with previous reports (Gruters et al., 2018; Lovich et al., 2023b; Sotero Silva, Kayser and Bröhl; 2025; Sotero Silva and Kayser, 2026). Prominently, the signals within each ear exhibited the typical opposition pattern for left- and rightwards saccades and comprised multiple peaks in the 50 ms following the saccade onset. A direct within-participant statistical contrast between ipsi- and contralateral saccades revealed multiple time windows (‘clusters’) of significant differences between directions (black bars, Fig. 1A, B; left ear: 8 - 17 ms, p=0.0184, D = 1.22; 20 - 34 ms, p = 0.0005, D = 1.35; and 80 – 92 ms, p = 0.0332, D = 0.68; right ear: 7 - 17 ms, p = 0.0014, D = 0.89; and 21 - 36 ms, p = 0.0003, D = 1.77). This stereotypical pattern of EMREOs was consistently observed in most individual participants, as reflected by confidence intervals in the group data and the individual participant data (Fig. 1A, B insets).

We also analyzed the saccade properties for the included epochs (Fig. 1C). This confirmed that the saccades were well aligned with the instructed saccade directions (average saccade angles: -178.2 ± 25.3 and 2.54 ± 25.7 deg, circular mean ± circular SD) and the saccade amplitudes did not differ between left- and right-wards saccades (paired t-test: p = 0.390, t = 0.86, D = 0.19, bf10 = 0.32).

### Vertical EMREOs

The EMREOs time courses for vertical saccades revealed a different profile compared to those for horizontal saccades (Fig. 2A, B). For upwards saccades the signal time courses were similar in both ears, with a systematically fluctuating time course. For downwards saccades, in contrast, the group-level time course revealed much less modulation and no pronounced peaks. A statistical contrast between directions revealed significant differences between saccade directions in both ears (left ear: 35 - 47 ms, p = 0.0025, D = 1.42; right ear: 35 - 47 ms, p = 0.0387, D = 0.88; and 70 - 86 ms, p = 0.0078, D = 0.77). The single participant data corroborated the observation that the vertical EMREOs are less consistent across participants (Fig. 2A, B insets). In particular for downwards saccades the data reveal very little consistency in the signal time course between participants, which is commensurate with a relatively flat group-average.

**Figure 2:**
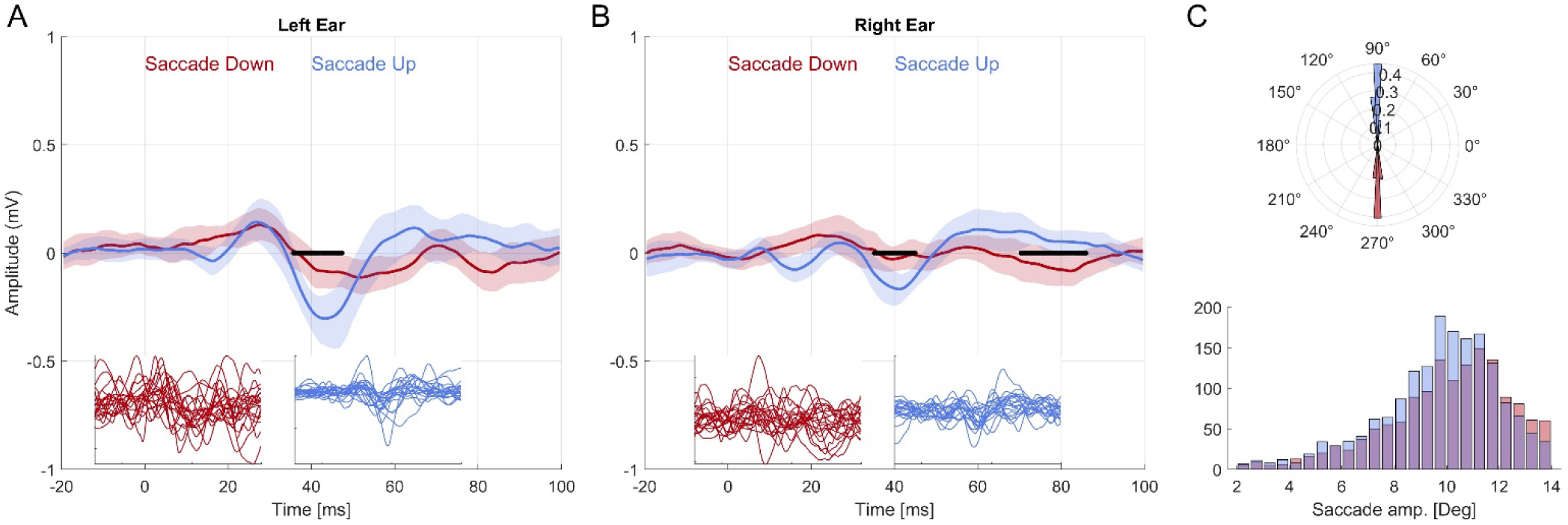
EMREOs induced by vertical saccades. **A, B)** Group-level EMREO time course (mean and 95% percentile bootstrap confidence interval across participants) for both saccade directions in each ear. Black lines indicate time windows with significant differences (cluster-based permutation test, at least p < 0.05). Insets show the individual participant EMREOs. For comparison the amplitude range on the axis is the same as for the horizontal data. **C)** Distribution of saccade directions (upper) and saccade amplitudes (lower) across all included saccades and directions.

As the EMREOs for downwards saccades revealed little modulation, we also performed statistical tests contrasting the group-level EMREOs to zero: hence testing whether there is a significant modulation in the signal relative to no modulation at all. For upwards saccades this revealed multiple significant clusters (left ear: 34 - 40 ms p = 0.0008, D = 1.33; right ear: 55 - 66 ms p = 0.0438, D = 0.57; and 35 - 46 ms, p = 0.0119, D = -1.10). The same was true for downwards saccades in the left but not the right ear (left ear: 16 - 32 ms, p = 0.0047, D = 0.77; 43 - 62 ms, p = 0.00570, D = -0.62; right ear: no significant clusters). This corroborates the notion that both upwards and downwards saccades induce an EMREO signal and that these differ from each other.

Finally, we directly tested whether each of the two vertical EMREOs differ from each of the two horizontal EMREOs. Since the main focus was a comparison of saccade directions we combined the p-values of both ears and different significant clusters of time points using the harmonic p-value. This showed that downwards EMREOs differed from left (p = 0.0287) and rightwards EMREOs (p = 0.0387), and upwards EMREOs differed from left (p = 0.0133) and rightwards EMREOS (p = 0.0308). Overall, this corroborates that the EMREOs induced by saccades in the four cardinal directions differ from each other.

Analyzing the saccade properties confirmed that the saccades followed the instructed directions well (Fig; 2C; average saccade angles: -92.7 ± 12.7 and 93.8 ± 23.3 deg). As for horizontal saccades, the saccade amplitudes did not differ significantly between directions (10.03± 0.29 vs 10.58 ± 0.23 deg; paired t-test p = 0.066, t = 1.95, D = 0.45, bf10 = 1.13).

### Reading out saccade vectors from EMREOs

Previous work has suggested that EMREOs provide parametric information about the saccade vector and allow telling, to some degree, the saccade direction from knowledge of the EMREO (Lovich et al., 2023b). However, this work also suggested that the information carried by EMREOs about the horizontal saccade directions is higher compared to vertical saccade directions, as the ability to reconstruct the saccade vector was better along horizontal than vertical directions (Lovich et al 2023b). The above observation that vertical EMREOs are modulated less strongly and less consistently across participants indeed suggests that EMREOs are indeed more informative about horizontal than vertical saccade features.

We explicitly tested this hypothesis in two ways, using similar tools as used in computational studies on neural information processing. First, we used a decoding framework and asked how well we can predict the categorical saccade direction (left, right, up, or down) from the combined single-trial EMREOs in the two ears. That is, we asked whether seeing the two EMREOs in a single trial, one can in principle tell the categorical saccade direction. In a second analysis we used a reconstruction framework and asked how well we can predict the precise saccade vector from single trial EMREOs. This analysis effectively asked whether single trial EMREOs provide direct parametric (i.e., continuous) information about the saccade vector. In both analyses we concatenated the EMREO time courses from both ears along the feature dimension, allowing the respective analysis algorithm to rely on whatever feature in the two signals carries the most relevant information. Both analyses converged to the same result: that EMREOs provide more information about horizontal than vertical saccade components and that the overall ability to faithfully reconstruct the two-dimensional saccade vector from EMREOs is only moderate.

The multivariate classifier was trained within each participant, hence allowing for idiosyncratic differences in the EMREO signals between participants. The overall classification accuracy was above chance (Fig. 3; 38.6 ± 1.5% correct, mean ± sem across participants, chance = 25%; paired t-test p < 10^-5^, t = 8.9). However, the confusion matrix associated with the classifier revealed a clear difference between horizontal and vertical dimensions: the percentage of correctly classified horizontal saccades (44.2 ± 2.4 % correct) was significantly higher compared to the percentage of correctly classified vertical saccades (32.9 ± 1.2% correct; paired t-test: p = 0.0001, t = 5.2, bf10 = 1233). Hence, the saccade-related information was consistently higher along the horizontal axis, but overall classification accuracy was also moderate.

**Figure 3:**
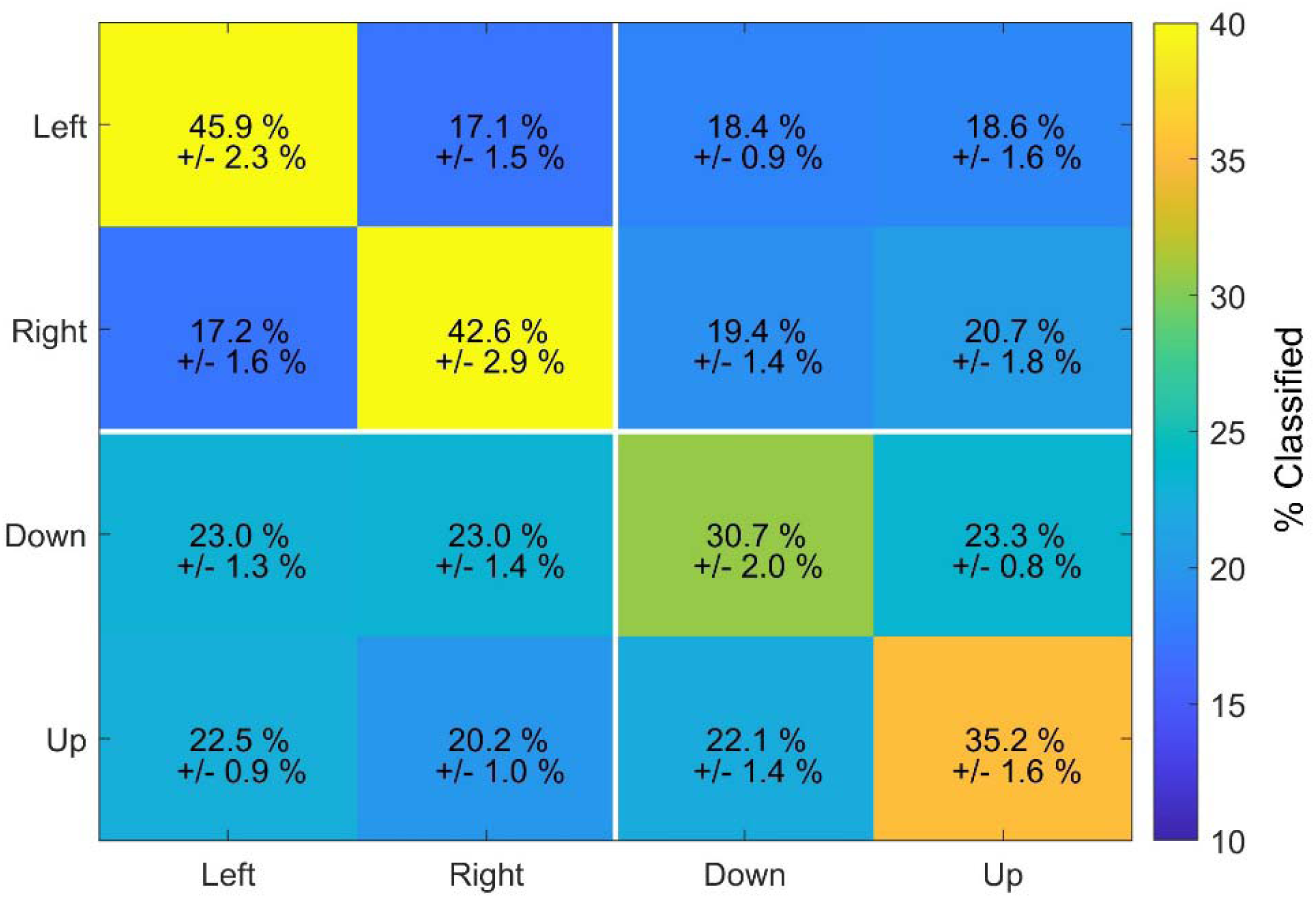
Classification of saccade direction based on binaural EMREOs. For each participant we classified the actual saccade direction based on the binaural single-trial EMREO epochs. The matrix shows how often (in %) each actual saccade direction was classified as each candidate direction (i.e., as a confusion matrix). Numbers indicate the average (and sem) across participants. Diagonal entries are correctly classified saccades, off-diagonal entries falsely classified saccades.

For the reconstruction analysis we used a regularized regression to predict the continuous horizontal and vertical saccade components based on the EMREO time courses.

The results in Fig. 4 show that the reconstruction was able to recover some of the spatial information in the saccade vector but vastly underestimated the saccade amplitude. At the group-level we tested the reconstruction accuracy against a null hypothesis of chance performance (reconstructing saccades at the screen center, 0 deg) using a 99% percentile bootstrap confidence interval. These are shown in Fig. 4 and the respective inset, demonstrating that the group-level reconstruction was significant for each direction (p-values based on the bootstrap for the null hypothesis of reconstruction of zero degrees: right p <10^-5^, left p < 10^-5^, up p = 0.003, down p = 0.018). To analyze the reconstruction performance in more detail, we calculated for each participant the correlation between the actual and reconstructed horizontal (vertical) saccade amplitudes. These correlations were significantly higher for horizontal compared to vertical saccade amplitudes (r = 0.41± 0.03 vs. r = 0.23 ± 0.02, paired t-test p < 10^-5^, t = 6.02, bf10 = 2569). We also calculated the root-mean-squared errors of the reconstruction: these were significantly smaller for horizontal compared to vertical saccades (rmse: 6.02 ± 0.11 deg vs 8.29 ± 0.17 deg; p < 10^-5^, t = 11.45, bf10 > 10^5^). Hence, the reconstruction ability was consistently higher along the horizontal dimension.

**Figure 4:**
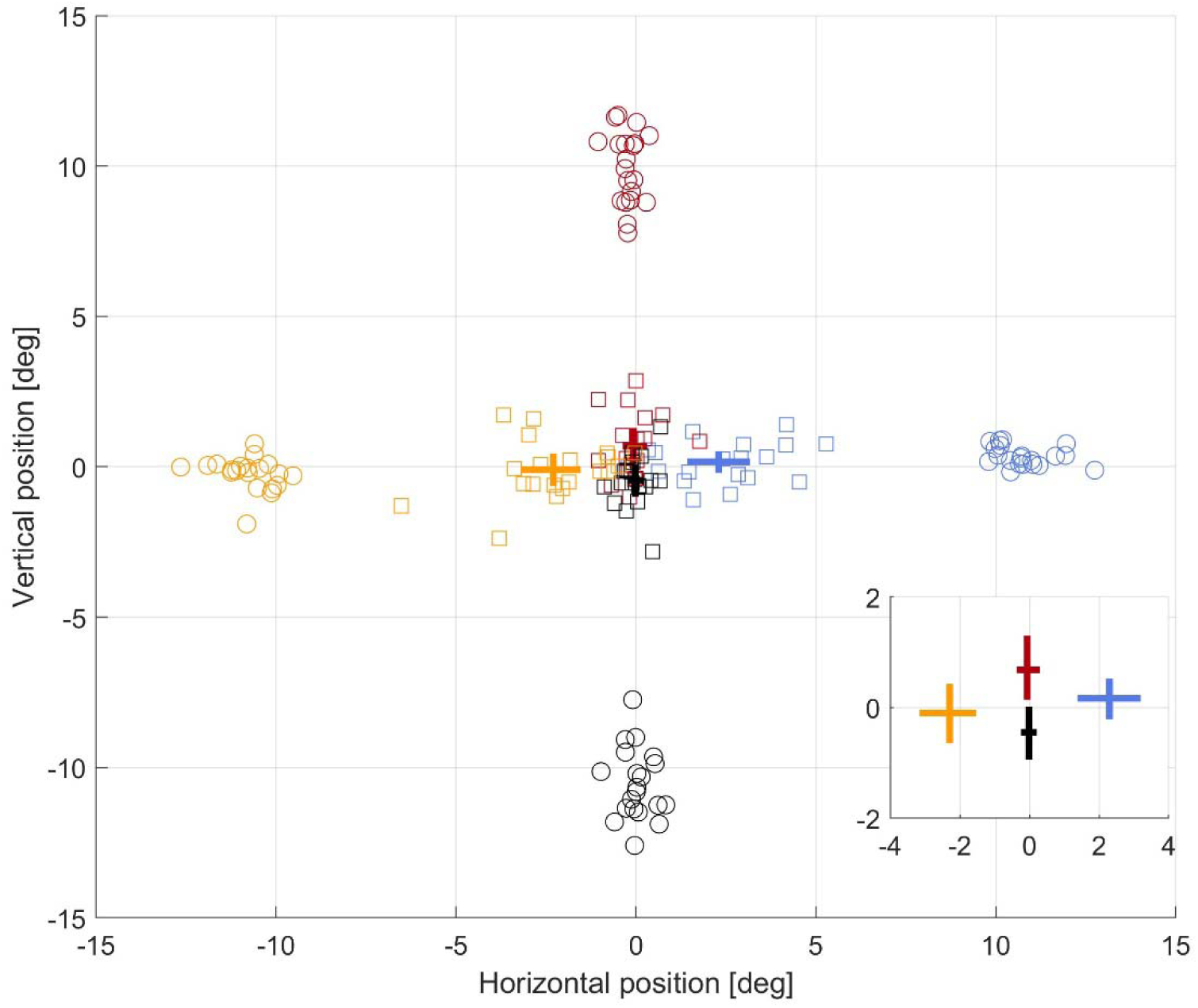
Reconstruction of saccade vectors based on binaural EMREOs. For each participant, we trained a reconstruction model predicting the actual saccade direction and amplitude based on the binaural single trial EMREO epochs. The figure shows the participant-wise averaged positions of actual saccades (color-coded by the actual saccade target) as circles and the respectively reconstructed average positions per participant as squares. The error-bars indicate the group-average and 99% percentile bootstrap confidence intervals for the horizontal and vertical saccade components. The inset enlarges the central part of the saccade range to visualize the confidence intervals around zero (screen center).

However, along both dimensions the model vastly underestimated the actual saccade amplitude showing that the direction of a saccade is much better recovered than the actual saccade size. Importantly, this underestimation cannot easily be overcome by simply scaling the model output, as at the group-level the variability in the reconstructed mean positions was on the same order of magnitude as the position itself. This is visible in the wide confidence intervals in Fig. 4, which in particular for the vertical dimension include the screen center. It is also reflected in the coefficient of variation (CV; standard deviation relative to mean) of the actual and reconstructed saccade amplitudes. In the actual data, the group-level CV of the actual mean saccade amplitudes were 0.067 and 0.097 for horizontal and vertical saccades, reflecting a small scatter around the average executed saccade amplitude. For the reconstructed data, the CVs were about ten times higher (0.69 and 0.76), reflecting a wide scatter around the average. Hence, the variability in the reconstructed saccade amplitudes precludes a reliable estimate of the average saccade amplitude, suggesting that based on this reconstruction model one cannot well predict the actual saccade vector based on the single trial EMREOs.

## Discussion

The EMREO signals have been studied in a growing number of experimental paradigms and contexts, yet both the origin and functional implications of EMREOs remain unclear (Gruters et al., 2018; Bröhl and Kayser, 2023; Lovich et al; 2023b; Abbasi et al., 2025; Leon et al., 2026; Sotero Silva, Bröhl and Kayser, 2026). One intriguing notion holds that if the underlying processes inducing the EMREO in the ear are indeed reflecting the precise saccade direction and amplitude, this may allow including spatially specific information about oculomotor behavior directly when encoding sounds in the ear. This, in turn, could help to combine the spatial signals received by the eyes and the ears to combine their respective spatial information about the location of objects in our environment (Razavi, O’Neill, and Paige, 2007; Recanzone, 2009; Cui et al., 2010; Krüger et al., 2016). Such a function would require that the EMREO signal carries reliable and precise information about saccade direction and amplitude on a trial-by-trial basis. However, whether this information is indeed available in single trial EMREO signals remains debated.

### EMREOs for horizontal and vertical saccades

Many studies have focused on saccades along the horizontal dimension, and the collective insight across these is that the EMREO signal induced by horizontal saccades is very reliable, consistent across participants and clearly related to the saccade direction: that is, the sign, amplitude and timing of distinct deflections in the EMREO depend on whether the saccade is made to a direction ipsi- or contralateral to the ear studied (Gruters et al., 2018; Bröhl and Kayser, 2023; King et al., 2023; Lovich et al., 2023b; Sotero Silva, Kayser and Bröhl, 2025). In contrast, whether horizontal EMREOs are also related to the amplitude of a saccade seems less clear. A number of studies have analyzed the relation between the EMREO signal and horizontal saccades using regression models (Bröhl and Kayser, 2023; King et al, 2023; Lovich et al., 2023a; Sotero Silva and Kayser, 2026). These revealed that the signal amplitude (and sign) at different latencies relative to saccade onset scale in proportion with the saccade property. However, these regression analyses tend to conflate saccade direction with saccade amplitude, leaving it unclear how much variability in the EMREO is directly explained by saccade amplitude irrespective of the direction itself (left vs right). While in the earliest reports of EMREOS a scaling of signal with saccade amplitude is visible (Gruters et al., 2023) a later study has directly quantified the relation between the amount of EMREO signal variation in the first 35 ms following saccade onset. This found that there was only a small increase in EMREO amplitude with saccade size and noted that the variation in EMREO with saccade amplitude was comparatively small (King et al., 2023).

Fewer studies have examined vertical saccades in detail. One study noted that for vertical saccades no consistent pattern in the EMREO waveform amplitude or latency could be discerned across participants (King et al., 2023). Another study reported that EMREOs induced by upwards saccades were larger than those for downwards saccades, similar to what we found here (Lovich et al., 2023b). That study also noted that vertical EMREOs revealed a less obvious counter-phase pattern than horizontal EMREOs. Our data corroborate these results and confirm that the EMREOs induced by vertical saccades differ from those induced by horizontal saccades and that vertical EMREOs by themselves are less reliable across participants. The single participant data here and in previous work show that the vertical EMREOs display large inter-individual variability, resulting in a much less stereotyped group- average signal.

The neural pathways and structures in the ear driving EMREOS remains unclear. Together with previous evidence our results suggest that either those pathways and structures giving rise to horizontal and vertical EMREOs are (partially) distinct, or that the temporal pattern in which these are activated for horizontal and vertical EMREOs differ. Prominently, horizontal saccades involve the conjugate association of antagonistic muscles across the eyes, creating strong left-right asymmetry in the engaged nerves (CN III and CN VI) and brainstem centers. In contrast, vertical saccades involve a more symmetric and bilateral activation of muscle pairs and cranial nerves (CN III) across the eyes. One possibility is that EMREOs primarily reflect this differential engagement of neural pathways. This would directly explain the largely opposing pattern for EMREOs along the horizontal axis and could also explain the smaller and less differentiated EMREO signal along the vertical axis, in particular for downwards saccades. Interestingly, the distribution of spontaneous saccades is also not uniform across the full 360° range, perhaps reflecting the statistical regularities of natural scenes or differences in oculomotor control (Carrasco et al, 2001; Foulsham 2008; Leigh and Zee, 2015). For humans, saccades along the cardinal directions are more frequent than along oblique directions, horizontal saccades more frequent than vertical ones, and upwards saccades are more frequent than downwards saccades (Foulsham 2008; Tatler and Vincent, 2009; Reeves et al., 2022). By virtue of EMREOs, this difference in oculomotor behavior is also reflected in the ear.

Still, the precise origin of what causes the EMREO signal remains unclear. Previous studies have associated this with the coordinated contraction of the tensor tympani and stapedius, and a possible contribution from outer hair cells (King et al., 2023; Lovich et al., 2023b; Sotero Silva, Kayser and Bröhl, 2025). These could either modulate reverberations of the eardrum or directly drive those. Interestingly, the two middle ear muscles are innervated by different cranial nerves (CN V for the tensor tympani and VII for the stapedius), though we lack precise knowledge about projections from oculomotor pathways to the middle ear muscles. In support of this, previous studies have noted that stapedius motor neurons are responsive to non-acoustic sensory signals (Salomon and Starr, 1963; Mukerji et al., 2010; Deiters et al., 2019; Tasko et al., 2022). However, it also remains unclear whether the tympanic membrane indeed moves directly in proportion to the acoustic EMREO signal recorded in the ear canal. One possibility is that the tympanic membrane only moves very briefly, or changes its tension, and the resulting temporally extended signal recorded is only an acoustic reverberation resulting from this brief change (Sotero Silva, Bröhl and Kayser, 2026). Measurements using other modalities than in-ear microphone recordings are required to further elucidate both the underlying neural pathways and actual physical changes in the ear during a saccade.

### Weak parametric information in EMREOs

We used two approaches to test whether and how much parametric information the bilateral EMREO signals carry about the inducing saccade. This is a critical test of the hypothesis that the EMREO could reflect an underlying mechanism that plays potentially a role in coordinating the encoding of visual and acoustic spatial signals. These are initially received in retinal-centered and head-centered coordinates and effective combination of multisensory spatial information requires the merging of information from different coordinate systems. If single-trial EMREOs allow recovering parametric information about saccades, this would show that at least such a role in multisensory coordinate alignment is possible in principle.

The classification analysis showed that the EMREO signals indeed hold significant information about the cardinal saccade direction. Yet, this information was significantly higher along the horizontal axis, in line with the horizontal EMREOs providing a richer and more reliable time course. The reconstruction analysis corroborates this, but also shows that estimating the precise saccade vector, including its amplitude, based on EMREOs is difficult. While the general direction was reconstructed well, the actual saccade amplitude was vastly underestimated, and the variability (uncertainty) in the reconstructed saccade amplitude was about as large as the average reconstructed amplitude itself. As a result, the underestimation of the reconstructed saccade amplitude could also not easily be overcome by scaling the reconstruction output, as this would exacerbate the uncertainty to the same degree. This finding corroborates a previous attempt to recover parametric saccade information from EMREOs (Lovich et al., 2023b): also, this study found that horizontal saccade components are recovered better than vertical ones, reported a severe underestimation of the saccade amplitude, and an average uncertainty in the reconstructed amplitude corresponding to about half the tested saccade range. Collectively, these results show that EMREOs indeed feature parametric information about saccade vectors but raise concerns about whether this information is sufficiently reliable to reflect a neural mechanism that could indeed contribute to the reliable alignment of audio-visual spatial information.

## Author contributions

NSS and CK Designed Research; NSS Performed Research; NSS and CK Analyzed data and wrote the paper.

